# Cortex deviates from criticality during action and deep sleep: a temporal renormalization group approach

**DOI:** 10.1101/2024.05.29.596499

**Authors:** J. Samuel Sooter, Antonio J. Fontenele, Cheng Ly, Andrea K. Barreiro, Woodrow L. Shew

## Abstract

The hypothesis that the brain operates near criticality explains observations of complex, often scale-invariant, neural activity. However, the brain is not static, its dynamical state varies depending on what an organism is doing. Neurons often become more synchronized (ordered) during unconsciousness and more desynchronized (disordered) in highly active awake conditions. Are all these states equidistant from criticality; if not, which is closest? The fundamental physics of how systems behave near criticality came from renormalization group (RG) theory, but RG for neural systems remains largely undeveloped. Here we developed a temporal RG (tRG) theory for analysis of typical neuroscience data. We mathematically identified multiple types of criticality (tRG fixed points) and developed tRG-driven data analytic methods to assess proximity to each fixed point based on relatively short time series. Unlike traditional methods for studying criticality in neural systems, our tRG approach allows time-resolved measurements of distance from criticality in experiments at behaviorally relevant timescales. We apply our approach to recordings of spike activity in mouse visual cortex, showing that the relaxed, awake state is closest to criticality. When arousal shifts away from this state – either increasing in more active awake states or decreasing in deep sleep – cortical dynamics deviate from criticality.

## Temporal renormalization group for brain dynamics

Diverse brain functions require complex neuronal population dynamics, with a broad repertoire of time scales: from fleeting response to sensory input (∼10-100 milliseconds), to execution of complex body movement sequences (∼100-1000 milliseconds), to even longer times scales of working memory (∼1-10 seconds). While some timescales are, in part, imposed by external influences (e.g. sensory input), most brain functions require internally generated dynamics (e.g. motor control and working memory). How does the brain generate such a broad range of time scales? Statistical physics offers a parsimonious hypothesis; the ‘criticality hypothesis’^1–3^ posits that by operating near the critical point of a phase transition, the intrinsic dynamics of many interacting neurons manifest with temporal scale-invariance over a broad range of timescales. The closer to criticality the system operates, the broader the range of scale-invariant timescales, with the range diverging at criticality^4,5^. In line with this hypothesis, a rich history of studies examining “1/f” power spectra^6–8^ and long-range temporal correlations^9–11^ of measured brain activity suggests that the brain often generates temporally scale-invariant population activity.

Moreover, many studies have reported evidence for criticality based on statistics of neuronal avalanches^12–15^. These studies report phenomenological evidence in favor of criticality in the brain, in agreement with a variety of computational models^2,16^, but a more fundamental theory of criticality and how it impacts the timescales of brain dynamics has been lacking. This phenomenological approach to investigating brain criticality has been fruitful but leaves open multiple important questions. For instance, as we will show here, there are multiple distinct types of critical dynamics, each temporally scale-invariant; which type of critical dynamics manifest in the brain? Is the brain always near criticality, or does it deviate from criticality under some conditions? Precisely how close to criticality can the brain be, how far does it deviate?

In non-living systems, renormalization group (RG) theory is responsible for fundamental theoretical understanding of criticality^17–19^ and answered questions akin to those we raised above. What are the prospects of a successful RG theory in the context of neural systems? Several recent studies have taken initial steps in this direction^20–24^, but fall short of our goals here. RG has been developed for neural models^23,24^, elucidating critical exponents^24^ and computational benefits of criticality^23^, but these approaches were not aimed at analyzing experimental data. Two recent RG approaches^20,22^, developed with data analysis in mind, were motivated by block-spin renormalization group ideas like those pioneered by Kadanoff^25^. In one case^20^, a binary neuron model was fit to empirical data and then the behavior of the best fit model was examined under coarse graining. A limitation of this approach was that their model assumed nearest neighbor interactions on a two-dimensional lattice; this assumption is not unreasonable at large spatial scales of the brain but is severely violated in more typical recordings of local populations of neurons (say in a ∼1 mm^3^ volume). In another case^22^, a model-free, “phenomenological” renormalization group (pRG) approach examined statistical properties of measured brain dynamics at different levels of coarse-graining, seeking fixed points. In RG, a ‘fixed point’ refers to statistical properties that are invariant upon coarse graining. This pRG approach has been used to support the criticality hypothesis in monkeys^26^ and mice^22,27^.

For pRG, no model of the brain is needed, unlike other approaches^20,23,24^ and unlike traditional RG in physics where a model of the physical system is needed, e.g. the Ising Hamiltonian. From one point of view, it is an advantage of pRG to be model-free; indeed, the brain is not sufficiently well understood to write down an indisputable model.

However, this model-free approach comes with disadvantages. Without a model, it is not possible to fully characterize the fixed points that exist. Thus, it is impossible to know what kind of criticality one should be looking for; without such concrete predictions, the empirical discovery of a possible fixed point lacks the certainty afforded by matching a prediction.

For our goals here, the most important limitation of previous RG approaches for analyzing data^20,22^ is that they require long recordings, and therefore are not time-resolved.

Considering many previous observations, the brain seems to shift among different dynamical regimes, depending on the behavior of the organism. For instance, focused attention, heightened arousal, and increased body movement are associated with reduced correlations among neurons, sometimes called a desynchronized cortical state^28–30^. At the other extreme of arousal, deep sleep and anesthesia are associated with elevated synchrony^31,32^. Such behavior-dependent neural coordination motivated our hypothesis that proximity to criticality varies depending on behavior. To test this hypothesis, we developed a temporal renormalization group (tRG) theory that is model-based and provides a time-resolved assessment of proximity to criticality and which type of fixed point is closest.

Our tRG approach is based on the order *p* autoregressive (AR(p)) model, for which the activity *x*_*t*_ at time step *t* is

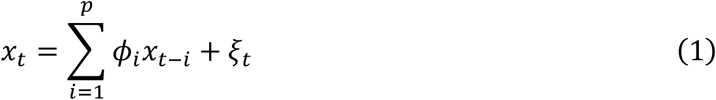

where *ξ*_*t*_ is independent gaussian white noise with variance *σ*^2^ and the history kernel *ϕ*_*i*_ determines how *x*_*t*_ is influenced by *p*time steps of recent history *i* = 1,2,.., *p*(Fig 1A). We chose the AR model for its simplicity and because it accurately reproduces the fluctuations traditionally used to study criticality in neural systems, as we will show quantitatively below. We first apply our theory to compare awake brain activity to that during deep (non-REM) sleep (Fig 1b,c). We re-analyzed spike data recorded from primary visual cortex and first reported by Senzai et al (2019)^33^. Activity fluctuations in the awake brain (Fig 1b) are qualitatively different from those during sleep (Fig 1c); the awake state is less synchronous and has more long timescale dynamics. Conceptually, our tRG theory describes how fluctuations change when viewed at different timescales, i.e. with different degrees of temporal coarse-graining, as illustrated in Figs 1d-f. For an AR model tuned precisely to criticality, the fluctuations are scale-invariant; they are statistically self-similar across different timescales (Fig d). Here, the coarse-graining was implemented by low-pass filtering and then rescaling time and amplitude. The awake mouse (Fig 1e) shows complex fluctuations over a 100-fold change in timescales, but is not precisely scale-invariant and, therefore, we should conclude it is not precisely at criticality. However, complex fluctuations in the awake mouse persist over a greater range of timescales compared to the example from the sleeping mouse (Fig 1f), which approaches white noise after a few steps of coarse-graining. The greater range of timescales in the awake case suggests that it could be closer to criticality than the sleep example, but how can we make this qualitative speculation more quantitative and concrete?

**Fig 1.**
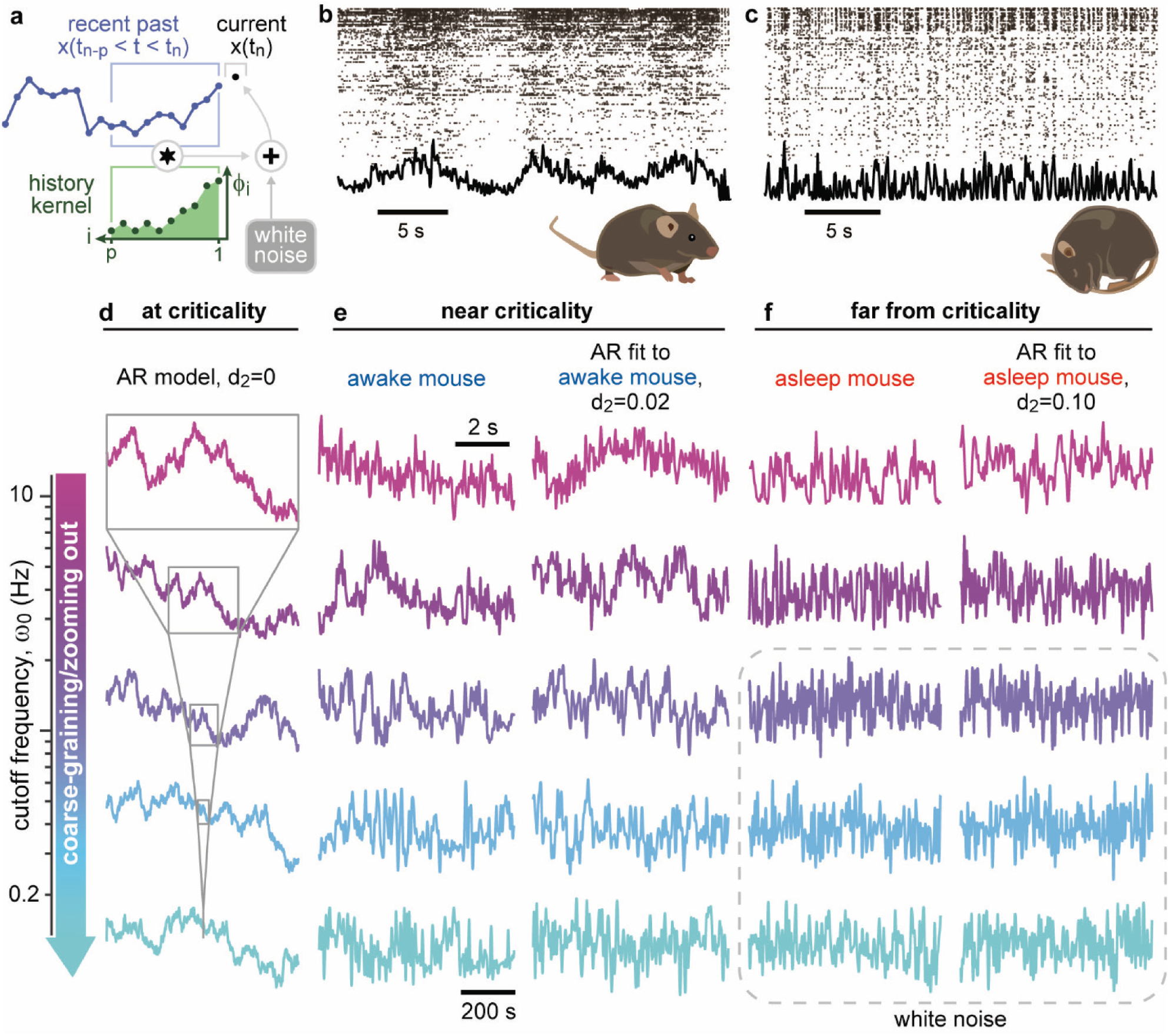
Coarse-graining suggests that brain dynamics are more scale invariant during wakefulness than sleep. **a)** Schematic cartoon of an order *p* autoregressive model AR(*p*). The history kernel *ϕ*_*i*_ is convolved with recent past activity and noise is added to generate the current activity. **b)** Example data from awake state, showing spike raster (top) and population activity (spike count time series, bottom). Note the mix of long and short timescale fluctuations. **c)** Example data from deep sleep. Note the lack of long timescale fluctuations. **d)** Example time series generated by an AR model at criticality. The fluctuations are temporally scale-invariant; they look the same when coarse-grained and rescaled in time and amplitude. **e)** Example population activity from awake mouse (left) exhibits complex fluctuations across 100-change of temporal scale. The best fit AR model exhibits similar multi-timescale fluctuations (right) and is close to the *β*=2 fixed point (*d*_*2*_=0.02). **f)** For the sleeping mouse, the population activity (left) and its best fit AR model (right) are further from the *β*=2 fixed point (*d*_*2*_=0.10) and quickly approach the white noise fixed point as they are coarse grained. The lesser degree of scale-invariance, as quantified by *d*_*2*_, suggests that the sleeping brain is further from criticality than the awake brain.

To go beyond such phenomenological coarse-graining, we employ our tRG theory of AR models, which is described with mathematical detail in Methods. The two main products of our tRG are 1) a theory-driven data analytic approach for assessing proximity to criticality, and 2) a roadmap of multiple kinds of scale-invariant dynamics (multiple tRG fixed points) in AR models. The first step of our data analytic approach is to fit an AR model to the time series in question (Methods). Next, we ask tRG, how close to criticality is the best fit AR model? Our theory provides precise, quantitative assessment of proximity to criticality for any given AR model (of arbitrary order), and for multiple different kinds of critical dynamics.

To be clear, neither the theory, nor the data analytic tools derived from the theory, implement phenomenological coarse-graining like that illustrated in Fig 1 and other RG approaches^20,22^. Indeed, one advantage of our approach is that subjective aspects of phenomenological coarse-graining are avoided (e.g. trying to judge when the activity has converged on or diverged from a fixed point). Rather, we mathematically determine how an AR model transforms into a new model under temporal coarse-graining, seeking fixed points of this flow in the space of models. For an AR(*p*) model there exist *p*+ 1 different fixed points (Supplementary Information). In the RG tradition, we interpret each fixed point as a type of criticality, except for the white-noise fixed point, which is scale invariant, but lacks the long-range temporal correlations expected for critical dynamics.

Whether or not any given AR(*p*) model flows to a fixed point is determined by the *p*coefficients *ϕ*_*i*_ of the history kernel. In the following, we index the fixed points with *β* = 0, 2, 4, …, 2*p*, where the power spectrum of the AR model has power-law form with exponent −*β* for low frequencies (the white noise fixed point has *β* = 0). Upon coarse graining, a family of AR models flow into each fixed point. The family for *β* = 0 is the largest, the family for *β* = 2 is second largest, and so on. By considering the *p*-dimensional space of all AR(*p*) models, we mathematically show that the geometry of the *p*+1 different families is rather neat and simple (Fig 2a, Supplementary Information). The family of *β* = 2 models occupies a (*p*-1) dimensional hyperplane, the *β* = 4 family occupies a (*p*-2) dimensional hyperplane within the *β* = 2 hyperplane, and so on. For any given AR(*p*) model, it is a straightforward geometrical calculation to measure the Euclidean distance from the model to each of these hyperplanes (along a line orthonormal to the hyperplane). We label these distances *d*_*β*_; the *d*_2_ reported in Fig 1e is the distance from the best fit AR model (order *p* = 20 in this example) to the *β* = 2 hyperplane.

**Fig 2.**
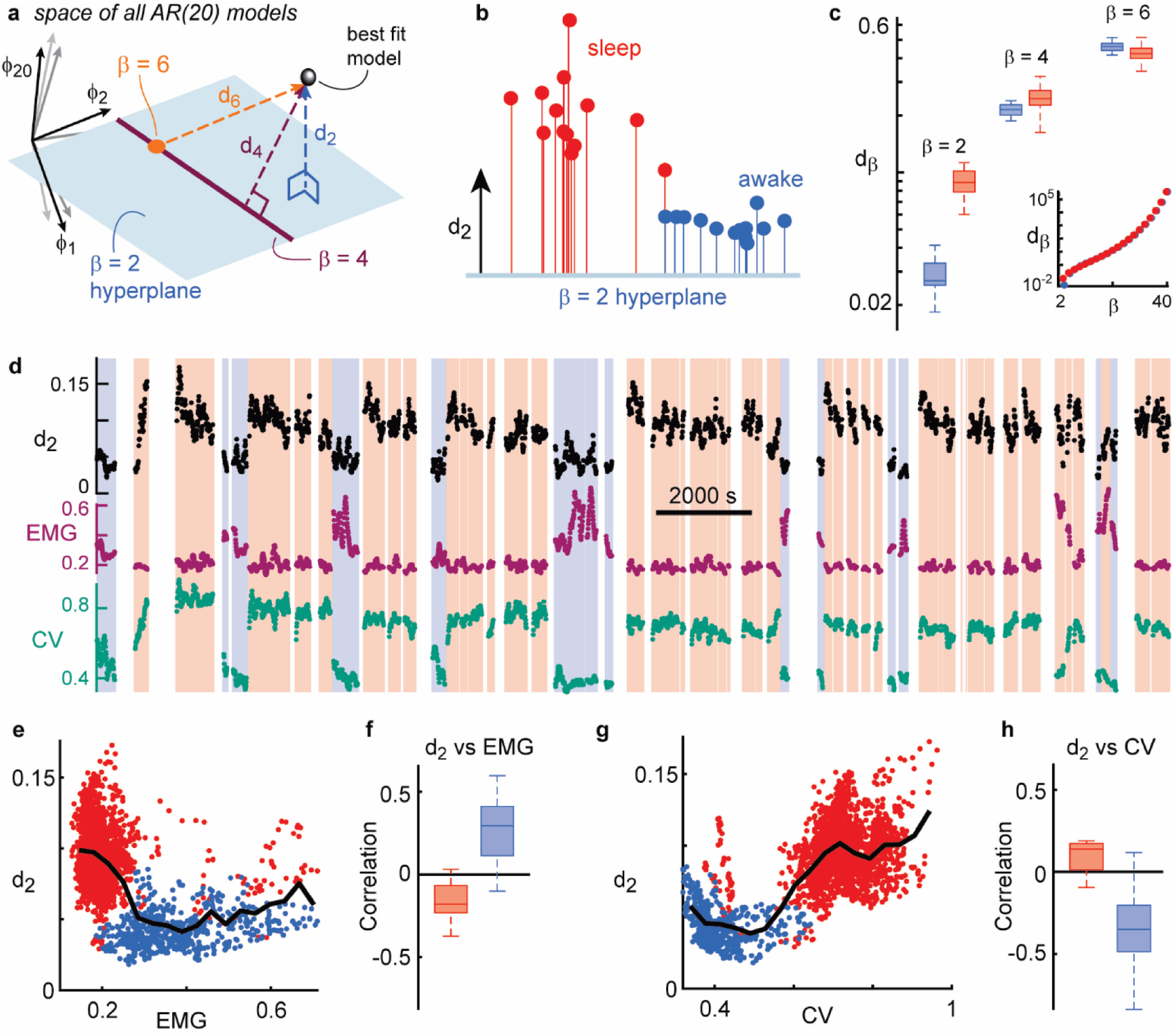
Deep sleep and awake bodily action drive deviations from criticality. **a)** Schematic cartoon of the space of all AR models and the nested structure of basins of attraction for the tRG fixed points. The distance *d*_*β*_ is the Euclidean distance between the best fit AR model (black point) and the nearest basin of attraction. **b)** For all 13 experiments, the best fit AR model for sleep (red) is further from criticality (*β*=2 type) than the best fit model for wakefulness (blue). **c)** Distances to other fixed points did not so clearly differentiate sleep versus wake. **d)** Computing *d*_*2*_ in a sliding time window (60 s duration) revealed time-resolved changes in distance to criticality consistent with the long-time summaries in panel b. Within the wake periods (blue background) *d*_*2*_ fluctuates together with body movement (EMG) and coefficient of variation (CV) of population activity. **e)** Example recoding showing that, during wake periods, *d*_*2*_ was correlated with EMG, indicating greater deviation from criticality for more intense body movement. Each point represents *d*_*2*_ and EMG from one 60 s window (red – sleep, blue – wake). Black line is a moving average of points. **f)** Summary of *d*_*2*_ vs. EMG correlations for all experiments. Box spans quartiles; line marks median; whiskers span range. **G)** For same example as panel e, *d*_*2*_ is anticorrelated with CV during wake. **h)** Summary of *d*_*2*_ vs. CV for all experiments.

### Deviations from criticality during sleep and awake action

In Fig 2b, we summarize 13 experiments (6.2±1.8 hours (mean±sd), 67±18 neurons per recording), comparing *d*_2_ for sleep versus wake periods. Fig 2b is a projection of the 20-dimensional space of AR(20) models, chosen such that the vertical direction of the plot is perpendicular to the *β* = 2 hyperplane and the horizontal direction is parallel to the *β* = 2 hyperplane. The brain dynamics in all 13 mice were closer to criticality during wake than during sleep.

The results comparing wake versus sleep (Fig 2b,c) were based on multi-hour recordings, but the first step of our data analytic approach – fitting an AR model to the data – does not require such long time series to obtain a good fit. In contrast, traditional methods of assessing proximity to criticality using spiking data (usually based on neuronal avalanche statistics) do require long time series. This opens an opportunity and question that has previously been unavailable: does the brain remain equidistant from criticality during the wake state or does proximity to criticality vary depending on behavior? To answer this question, we performed our measurement of *d*_*2*_ in a sliding 60 s window (with 5 s sliding increments). First, we found that such time-resolved *d*_*2*_ measurements were consistent with the measurements based on longer time series, exhibiting clear increases during deep sleep compared to wake (Fig 2d). Next, we compared fluctuations in *d*_*2*_ to measurements of muscle activity of the mice (electromyogram, EMG), which were recorded simultaneously with the brain activity. We found that *d*_*2*_ tended to increase when the mice became active (increased EMG, Fig 2e,f). We also compared *d*_*2*_ to the coefficient of variation (CV) of population spike activity (Fig 2g,h); decreased CV is associated with arousal^30^. Considering wake periods, we found that *d*_*2*_ was positively correlated with EMG (Spearman rank correlation, Fig 2f) and negatively correlated with CV (Fig 2h). During sleep, these correlations tended to reverse. Together these results demonstrate that cortex is closest to criticality (i.e. has lowest *d*_*2*_) in an awake, quiescent state. Elevated body movement causes *d*_*2*_ to increase. This result is qualitatively similar to previous studies that reported deviation from criticality in desynchronized and hyper-synchronized states in anesthetized animals^32^.

When using our tRG approach, two important parameter choices are the AR model order and the time bin used to create the population spike count timeseries; both can impact the quantitative values of *d*_*2*_. The primary results here (Fig 2) were robust over a range of model orders (5 to 20) and time bin choices (10 to 160 ms) (Supp Fig S1). Moreover, we verified that *d*_*2*_ can accurately track changes in dynamical state over a wide range of time scales for our choice of a 60 s sliding window duration (Supp Fig S2). Finally, as a control, we performed *d*_*2*_ analyses after removing correlations among neurons, but preserving all single-neuron spike statistics (shifting all spike times of each neuron by a random constant between 0 s and half the recording duration). We found that *d*_*2*_ = 0.75±0.05 for this control, which is much greater than our measurements of *d*_*2*_, even for sleep.

### Benchmarking against models and traditional methods

How does our tRG-based analysis compare to traditional methods for studying criticality in neural systems - neuronal avalanches, power spectra, and detrended fluctuation analysis (DFA)? We found that avalanche size and duration distributions, power spectra, and DFA fluctuation functions all exhibit power-law scaling that extends over a greater range of time scales during wake than during sleep (Fig 3). This observation agrees with our *d*_*2*_ results in Fig 2. Indeed, as a system deviates from criticality, it is expected to exhibit a decreasing range of power-law scaling (Fig 4). We also used these traditional statistical assessments to verify that AR models are, in fact, a good fit to the experimental data.

**Fig 3.**
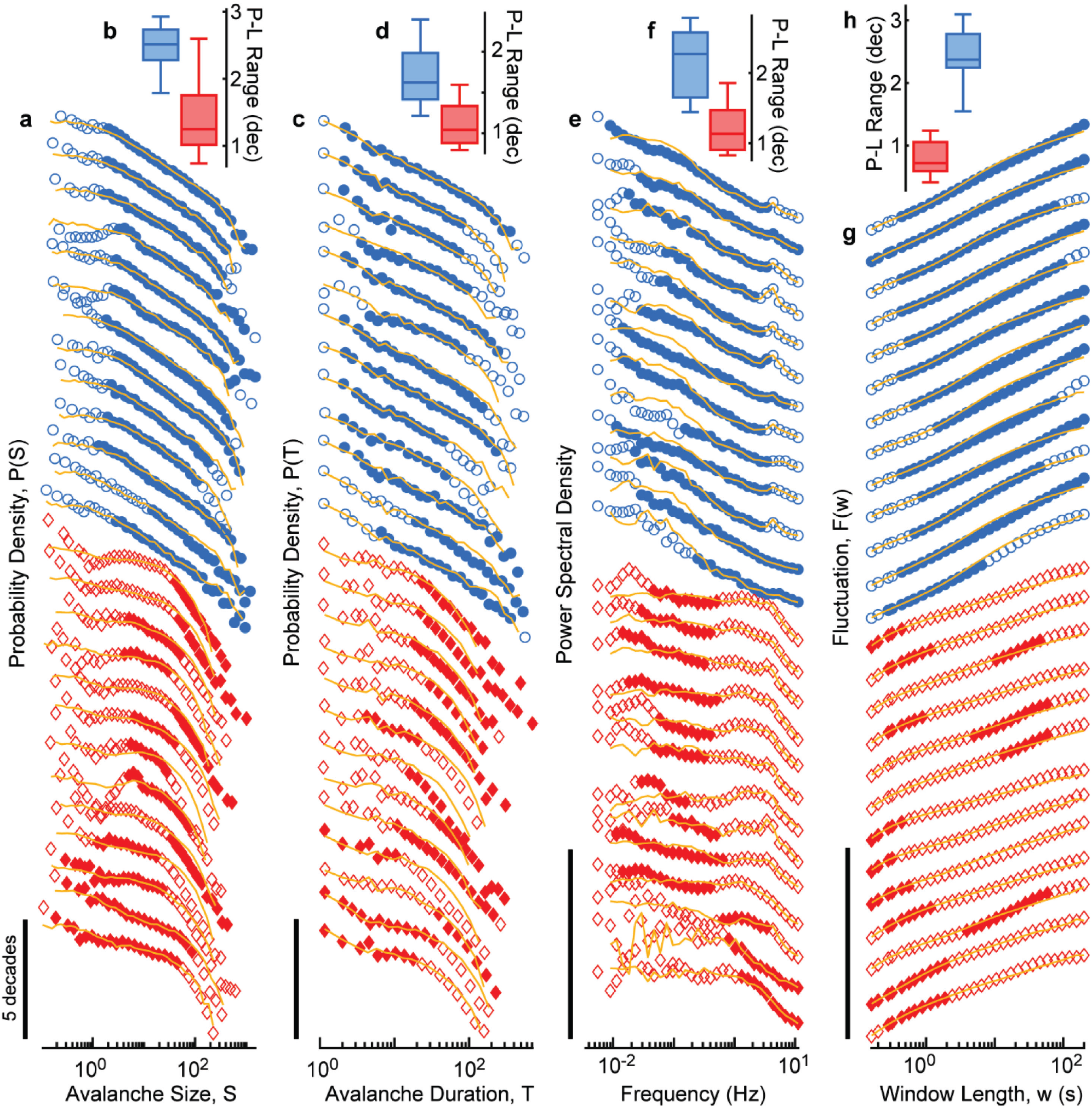
tRG approach agrees with traditional methods. **a**,**b)** Avalanche size distributions (for all recordings) during wake states (blue) exhibit a wider range of power-law scaling (P-L Range) compared to those during sleep (red). Distributions are shifted vertically for visual comparison. Solid points indicate the portion of the distribution with statistically significant power-law scaling. Yellow lines represent avalanches simulated with the corresponding best-fit AR model. **c**,**d)** Same as panels a and b, but for avalanche durations. **e**,**f)** Same as panels a and b, but for power spectral density. **g**,**h)** Same as panels a and b, but for DFA fluctuation functions.

**Fig 4.**
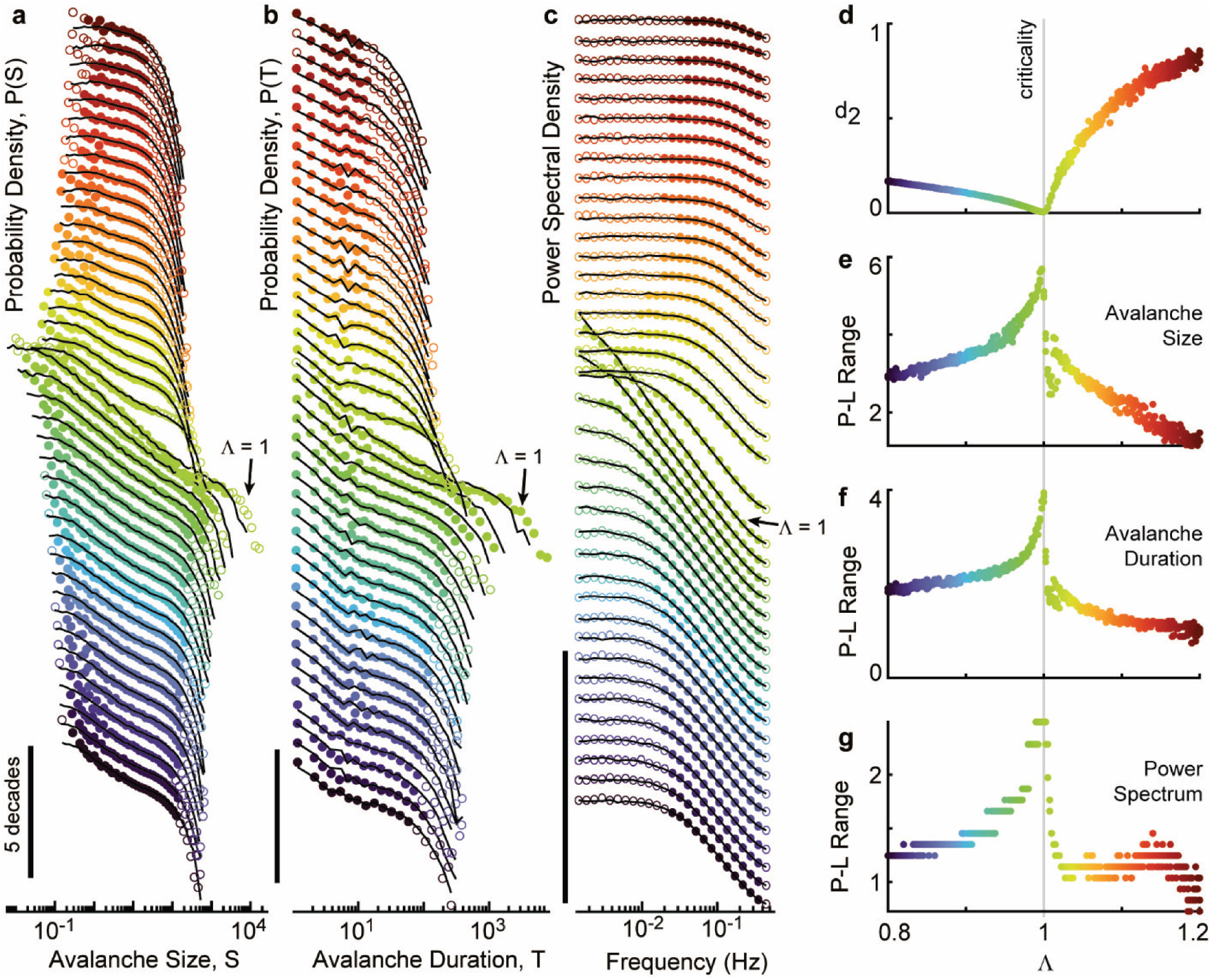
tRG approach accurately measures distance from criticality in ground truth model. **a)** Avalanche size distributions for a family of models ranging from subcritical (blue, Λ=0.8) to criticality (green, Λ=1.0), to supercritical (red, Λ=1.2). Black lines are simulated with the best fit AR model for each model Λ. Filled circles indicate the part of the distribution that is statistically well-fit by a power-law. **b)** Same as panel a, but for avalanche durations. **c)** Same as panel a, but for power spectra. **d)** *d*_*2*_ smoothly goes to zero as Λ approaches 1, thus reflecting indicating distance to criticality. **e-g)** Power-law range gradually decreases as Λ departs from 1.

After fitting an AR model for each experiment, we then used the best fit model to simulate a new time series. These simulated time series were generated in segments with the same durations as found in each experiment. We ran avalanche analysis and spectral analyses (power spectra and DFA) on these simulated time series and found very good agreement with those based on the actual data. Thus, basing our approach on AR models is well justified.

Next we benchmarked our approach against an established model for which ground truth knowledge of proximity to criticality is known^34,35^. The model consists of 1000 binary neurons (80% excitatory, 20% inhibitory, Methods). This model can be tuned through a range of states from an asynchronous, low firing rate phase, through a critical point, to a saturated high firing-rate phase. These states are conveniently parameterized by the largest eigenvalue Λ of the connectivity matrix. The critical point for this model is at Λ = 1. We draw three conclusions from the model (Fig 4). First, in the same way that we did for the experiments, we establish that AR models provide a good fit to the dynamics generated by the binary neuron model, accurately reproducing avalanche distributions, spectra, and DFA for a wide range of Λ (Fig 4a-c). This extends the validity of our approach. Second, and most importantly, the model confirms that *d*_*2*_ is very close to zero at criticality (at Λ = 1) and departs smoothly from zero as the model deviates from criticality (Fig 4d). Finally, the model verifies that power-law range gradually decreases as the system is tuned away from criticality (Fig 4e-g). Taken together, these results provide a ground truth example of how *d*_*2*_ measures the range of scale-invariant timescales and, thus, measures distance from criticality.

### Resolved controversy, limitations, future prospects

Previous claims about whether or not the brain operates close to criticality during sleep or active awake states are controversial. Some studies report deviation from criticality during sleep^5,36^, while others suggest the oppposite^31,37,38^ or little difference between sleep and wake^39,40^. Another related line of work shows that prolonged lack of sleep results in deviation from criticality, while sleep tunes the brain back to criticality^41,42^. For awake animals, one study showed in mice that the crackling noise scaling law for neuronal avalanches does not hold during running^43^, which is consistent with deviation from criticality. However, other avalanche-based studies suggested proximity to criticality in motor cortex does not change as a monkey executes a motor task^44^ or a mouse learns a motor task^45^. One possible reason for some of the discrepancies in the literature is that conclusions based on avalanche statistics often rely on estimates of power-law exponents from avalanche distributions, which are known to depend on subjective choices of analysis parameters^14,46^. We contend that our tRG-based approach provides more reliable estimates of proximity to criticality. Our approach directly quantifies temporal scale-invariance and thus establishes, without speculation, that the sleeping brain comes with less scale invariance than during wakefulness and that bodily action also reduces scale-invariance compared to the awake resting state. These differences in scale-invariance strongly suggest that both deep sleep and awake action cause deviations from criticality.

The two most important limitations of our theory and data analytic approach concern the kinds of data our approach should be applied to. First, our approach is best suited for recordings of neurons from a small volume (∼1 mm^3^) within which connectivity is approximately independent of spatial positions of neurons (more details in Supplementary Information). In contrast, our approach is not suited to spatially extended measurements like widefield imaging and functional magnetic resonance imaging. At these larger spatial scales, connectivity among neurons is more strongly dependent on distance between neurons and, thus would be better suited to an RG approach that accounts for space. We are optimistic that our approach can be generalized in future work to account for both spatial and temporal scale invariance. Second, not all neural systems generate population spike activity that is well-fit by an AR model. For example, in vitro preparations and some anesthetized states often generate bursty dynamics - quiet periods punctuated by population-wide volleys of spikes^12,32^. Such activity tends to have highly asymmetric distributions of activity, which are not well fit by AR models, and thus will require a different approach to assess criticality.

Nevertheless, our tRG approach offers several important advances compared to traditional studies of criticality in neural systems. Perhaps the biggest practical advantage is the time resolution of our tRG-driven data analytic tools. This time resolution allows more direct tests than before concerning the long-hypothesized functional implications of criticality.

Indeed, different behaviors come with different computational requirements; our approach may reveal how proximity to criticality shifts to accommodate these changing computational needs.

### Online Methods

*<u>tRG theory</u>* An AR model (Eq. (1)) can, in principle, generate any time series; the probability density for it to produce a given time series *x* ≔ {*x*_*t*_}_*t*_ is

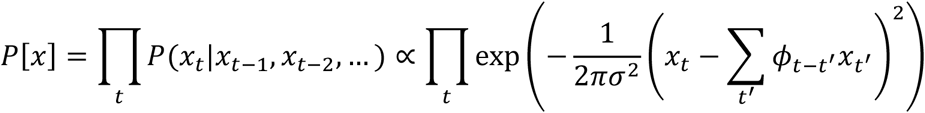

After some manipulations, we can rewrite this in terms of the Fourier transform 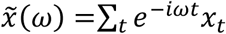 of the time series 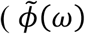 defined correspondingly) as

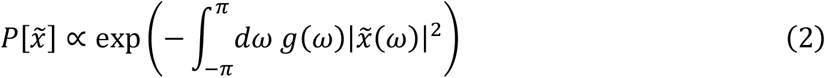

where

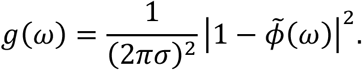

An equation of the form of Eq. (2) in fact holds for all stationary gaussian processes, with *g*(ω) proportional to the reciprocal of the power spectrum (Appendix A.2 of ref^47^).

This makes the coarse-graining step of the renormalization group—marginalizing over the Fourier components above a cutoff frequency *ω*_0_—straightforward. Formally, letting 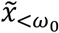 be the collection of Fourier components below the cutoff frequency *ω*_0_ and 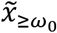 that above the cutoff, the distribution of trajectories factors as 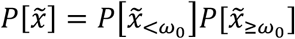 with

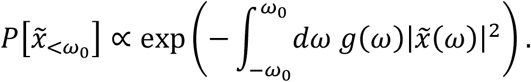

This differs from the original distribution in Eq. (2) only by the frequency bounds on the integral. Rescaling *ω* → *ω* ′ = *πω*/*ω*_0_ and 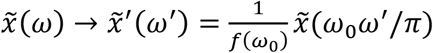 restores the original frequency bounds and, for an appropriately chosen rescaling function *f*(*ω*_0_), allows the distribution of the coarse-grained trajectories to flow into a fixed point as we take the cutoff frequency *ω*_0_ → 0. In terms of the rescaled variables,

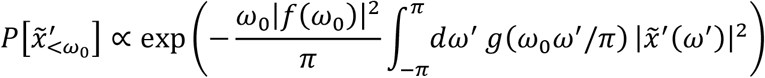

which has the same form as the original distribution with *g*(*ω*) replaced by

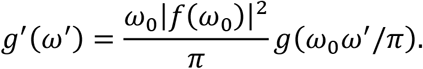

In the limit *ω*_0_ → 0, the dominant term in the Taylor series for *g*(*ω*) controls the behavior of *g*′ (*ω* ′). Since *g*(*ω*) is an even function of *ω*, its dominant term must be an even power of *ω*. For a given even *β* ≥ 2, we show in Supplementary Information that the dominant term in *g*(*ω*) is *ω*^*β*^ if and only if *a*_0_ = 1 and *a*_1_ = … = *a*_*β*/2−1_ = 0 ≠ *a*_*β*/2_, where *a*_*m*_ = ∑ _*t*_*t*^*m*^*ϕ*_*t*_ for all *m*. Setting the rescaling function to 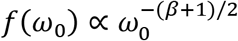 then makes *g′* (*ω* ′) → *ω* ′^*β*^ as *ω*_0_ → 0. Otherwise, the dominant term is the constant (*β* = 0) term and *g′* (*ω* ′) → *ω* ′^0^, i.e., the model flows into the white noise fixed point. We conclude that each value of *β* corresponds to a different renormalization group fixed point and that these are the only fixed points an AR model can flow into.

#### Calculating distances d_β_

There are four steps involved with calculating the d_β_ distances discussed in the main text. First, a spike count time series is created, which entails breaking time into consecutive time bins of duration Δt and then counting spikes, from the entire population of recorded neurons, in each time bin. Second, the spike count time series is z-scored (mean is subtracted and then it is normalized by standard deviation.) Third, an autoregressive model is fit to the z-scored spike count time series. For this we used the Yule Walker method^48^. Fourth, the coefficients of the best fit AR model history kernel are used to compute the distances *d*_*β*_ (Supplementary Information).

#### Avalanche analysis

We start with the same spike count time series used to calculate distances d_β_. Following established methods^14,49^, an avalanche is defined as an excursion of this time series above a fixed threshold, which we set to the 10^th^ quantile of the time series. The size of an avalanche is the area between the threshold and the time series over the course of the avalanche.

#### Spectral analyses

To calculate the power spectra in Figs. 3 and 4, we used Welch’s method with windows of length 200 s and 50% overlap between windows. The power spectra were resampled at 10 frequency points per decade, equally spaced logarithmically.

For the DFA fluctuation function, we followed the procedure in ref^11^. Briefly, for a given *w*, we split the z-scored spike-count time series into windows of length *ww* (50% overlap between windows), took the cumulative sum within each window, subtracted the least-squares fit line from this cumulative sum, and calculated the standard deviation of the resulting time series. The fluctuation *F*(*w*) is then defined as the average over windows of this standard deviation.

#### Handling segmented time series

Each experimental recording was broken into multiple relatively short segments to separate sleep epochs from wake epochs (e.g. Fig 2d). When performing AR model fits and calculating the avalanche distributions, power spectra, and fluctuation functions for non-REM and wakefulness, we accounted for the separation between different epochs of these states as follows:

1. For fitting AR models, part of the Yule-Walker method involves calculating the sample autocorrelation function. For this, we only considered within-epoch pairs of time points.
2. When defining avalanches, we excluded avalanches that temporally overlapped the start or end of an epoch.
3. In the spectral analyses, we only considered within-epoch time windows.

#### Power-law range

To calculate the power-law range of each avalanche distribution, we used the maximum-likelihood fitting algorithm described in ref^14^ with a goodness-of-fit criterion of 0.95 to calculate the range, in logarithmic decades, over which the avalanche size and duration distributions were well-fit by a power-law. To estimate the range of power-law scaling in the power spectrum and the DFA fluctuation function, we used a similar procedure. We systematically moved a lower cutoff *x*_*m*_ and an upper cutoff *x*_*M*_, performed a least-squares linear regression (in log-log coordinates) on the points between the cutoffs to get the best-fit power-law, and calculated the fraction of points within a band of width 2*ϵ* centered on that best-fit power-law. We call this the goodness-of-fit. The power-law range is then defined as the maximum value of log_10_(*x*_*M*_/*x*_*m*_) over pairs of cutoffs *x*_*m*_, *x*_*M*_ for which two conditions were met: (1) the goodness of fit exceeds 0.95 and (2) the exponent of the best-fit power-law is less than −0.3 (for the power spectrum) or greater than 0.8 (for DFA). The second condition excludes the case of white noise, which has a power-law power spectrum with exponent 0 and a power-law fluctuation function with exponent 0.5. We chose *ϵ* = 0.1 for the power spectrum and *ϵ* = 0.03 for the fluctuation function.

#### Binary spiking model

The results in Fig. 4 are based on the following model, previously studied in refs^34,35^. The model consists of *N* = 1000 binary spiking neurons; the *i*^th^ neuron spikes at time step *t* with probability

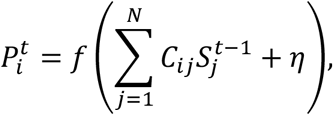

where *f* is the identity for arguments in [0, 1] and is capped at zero and one for arguments less than zero and greater than one, respectively. Here *C* is the connectivity matrix, 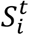 is unity if the *ii*th neuron spiked at time step *t* and zero otherwise, and η = 0.001 is the background spiking rate. Each entry in *C* is initially set to unity with probability 0.1 and to zero otherwise. 20% of the neurons are then selected at random to be inhibitory neurons and the corresponding columns of *C* are scaled by −1. Finally, we scaled every entry of *C* by the same positive constant to set the magnitude of the maximum eigenvalue of *C* to the desired value Λ. For each choice of Λ in Fig. 4, we ran the model three times, for 5 × 10^6^ time steps in each run.

#### Experimental data

The experimental data analyzed here was collected and first reported by Senzai et al (2019). The data is publicly available from the Buzsaki Lab data repository (https://buzsakilab.com/wp/database/). We analyzed 13 data sets with the following labels in the Buzsaki Lab database: YMV02, YMV07, YMV08, YMV09, YMV10, YMV11, YMV12, YMV13, YMV14, YMV15, YMV16, YMV17, YMV18. We excluded other recordings in this data set if they had less than 30 min total of deep sleep or less than 30 min of wakefulness. In brief, the data consisted of extracellular recordings (64-channel, linear, single shank silicon probe, Cambridge NeuroTech H3 64×1 probe) of single unit activity (67±18 units per recording) in mouse visual cortex (anteroposterior: +1.0 mm, mediolateral: +2.5 mm relative to bregma) spanning all cortical layers (20 μm inter-electrode spacing). Spike sorting was performed semi-automatically, using Kilosort^50^. Recordings were multiple hours (6.2±1.8), during which the mice naturally switched between sleeping and waking multiple times. Included in the data sets are classifications of wake, REM sleep, and nonREM sleep. These classifications were made by the Buzsaki Lab based on broadband local field potential (1-100 Hz), narrowband theta frequency LFP (5-10 Hz), and electromyogram (EMG), as described previously^33^.

## Data and code availability

All data is freely available from the Buzsaki Lab data repository (https://buzsakilab.com/wp/database/). The data analysis codes used to calculate *d*_*2*_, avalanche analyses, spectral analyses, and the network level binary neuron model are freely available on Figshare: 10.6084/m9.figshare.25927081

## Author contributions

JSS led mathematical and data analytic efforts. CL, AKB, and WLS contributed to mathematical efforts. AJF contributed data analysis. JSS, AJF, CL, AKB, and WLS conceptualized the study and acquired funding. JSS and WLS created figures and wrote the manuscript.

