## Supplementary Material for "Cortex deviates from criticality during action and deep sleep: a temporal renormalization group approach"

### Supplementary Information

#### Designing tRG for typical experimental data

Experimental study of critical phenomena in non-living systems has traditionally been based on measurements of macroscopic properties (e.g. magnetization); details at the microscopic scale (e.g. single molecules) are difficult to measure. In contrast, measurements in neural systems are commonly done and best understood at the smallest, most detailed scales of the system - single neurons and single action potentials (spikes). Thus, we aimed to develop RG specifically for application to typical recordings of mammalian brain activity with single-neuron resolution. Our theory and tools are best suited to high-density single-unit electrophysiology and calcium imaging of a local population, say 100 to 1000 neurons within a ~1 mm$^3$ volume. At this scale, the connections among neurons are weakly dependent on the spatial positions of the neurons[1–4], unlike physical systems with nearest-neighbor lattice interactions (e.g. Ising model). Since spatial positions play little role, we developed our RG to study scale-invariance in the time domain. Additional support for this approach is found in computational models with spatially-independent connectivity among neurons; in such models critical phenomena are temporal, not spatial[5–8].

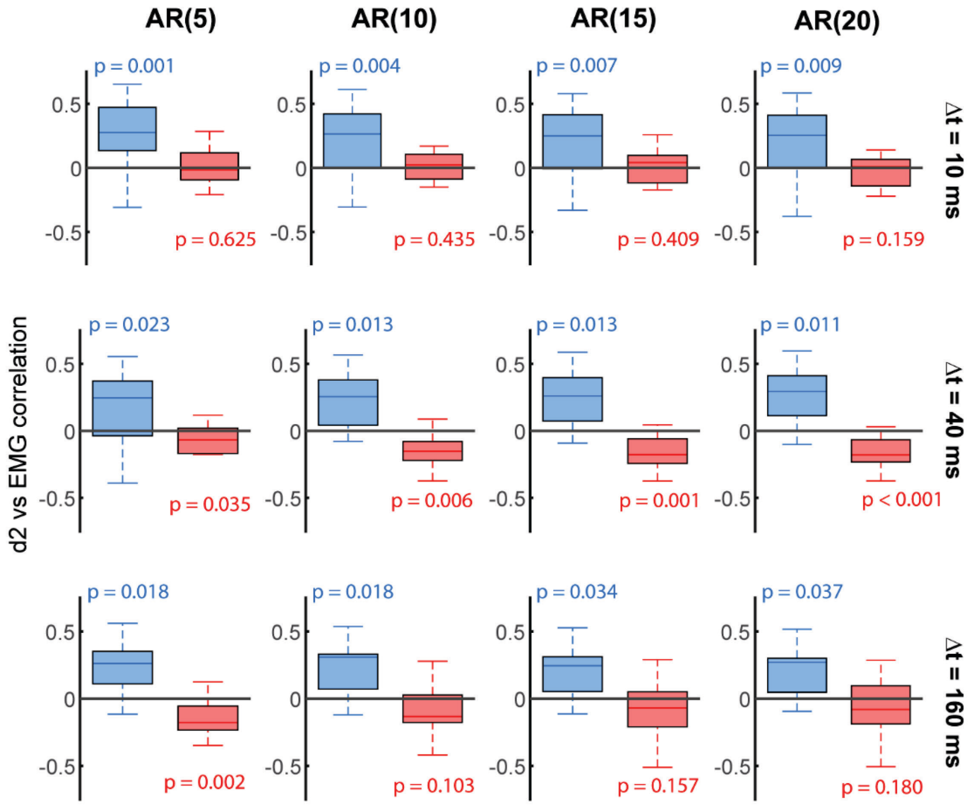


**Fig S1. Robustness to AR model order and time bin duration.** Here we show that the results in the main text (Fig 2f,h) are not very sensitive to the choice of AR model order, nor the choice of time bin (Δt) for creating the spike count time series. For AR model orders 5, 10, 15, 20 (columns, left to right) and Δt ranging from 10 ms to 160 ms (rows, top to bottom), the primary result that $d_2$ is positively correlated with EMG remains significant. The right column and middle row are the same parameters used in the main text. Here, p values are derived from a right-tail one-sample t-test for wake (blue), thus representing the probability of the null hypothesis that the correlation coefficients have zero mean versus positive mean. For sleep (red), p values are derived from a left-tail one-sample t-test, thus representing the probability of the null hypothesis that the correlation coefficients have zero mean versus negative mean.

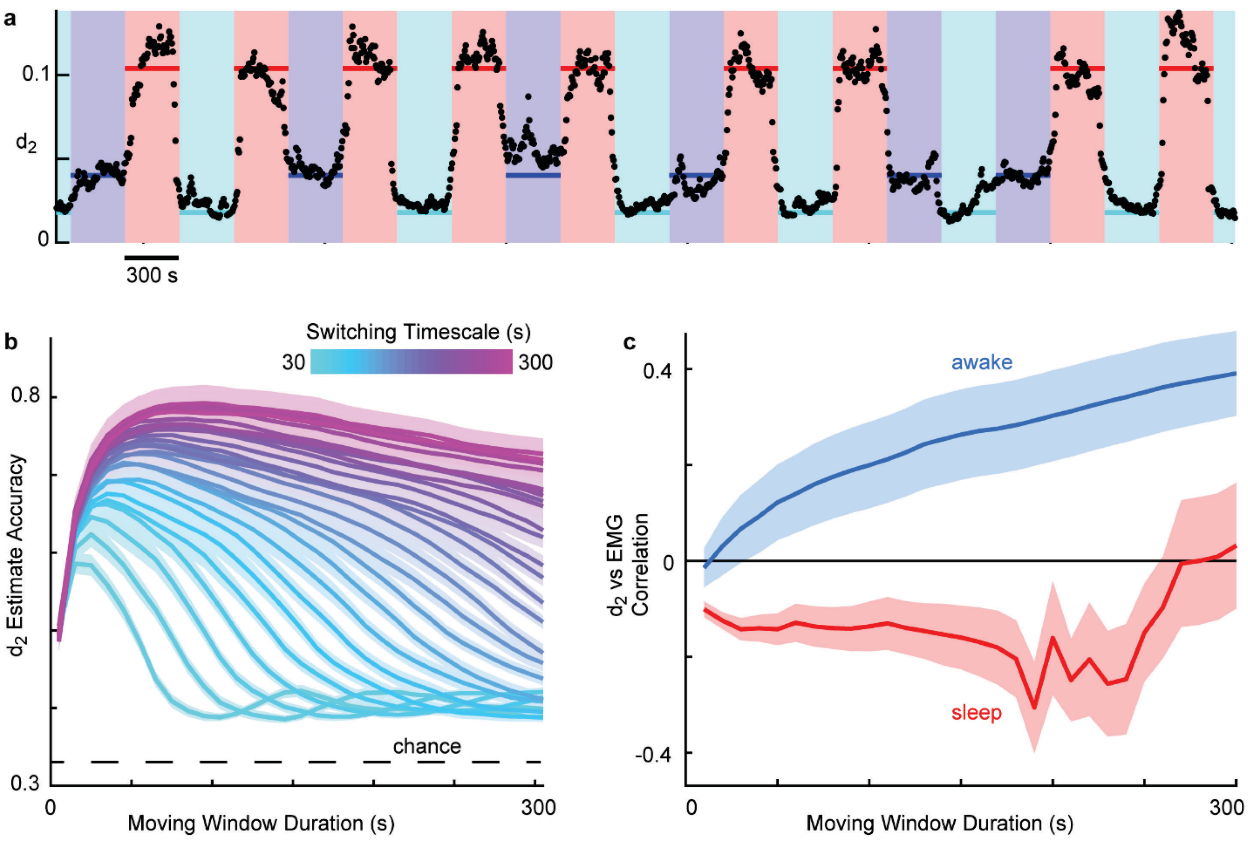


**Fig S2. Ground truth test of time resolution of $d_2$.** **a)** The black points represent $d_2$ estimates for a surrogate data set with controlled state transitions. Three states were simulated for $T = 300$ s each in randomized order (background color indicates state). The true $d_2$ for each state is indicated with the horizontal red, purple, and cyan lines. We used a 60 s sliding window with 55 s overlap. **b)** We systematically varied the switching timescale $T$ from 30 to 300 s (color) and examined the accuracy of $d_2$ for different sliding window durations. In general, there is a tradeoff: a very short sliding window duration precludes good accuracy due to poor AR fit (too few samples for a good fit), while a long sliding window that is longer than $T$ will lose accuracy due to mixing multiple states together. The shaded band behind each line represents mean±SEM, summarizing different choices for the three ground truth states – one based on each experiment. **c)** We found that the $d_2$ vs EMG correlations reported in Fig 2 was robust across a wide range of window durations.

#### Validating time resolution of $d_2$

The results in Fig 3 in the main text confirm that our measured $d_2$ differences between sleep and wake are consistent with traditional criticality measures, but it is not possible to use such traditional measures to back up our time-resolved $d_2$ results (Fig 2d-h), because these traditional methods require long time series for reliable results (especially avalanche analysis). In principle, any approach to measure time-resolved changes in state depends on two basic limitations. First, if the window in which we are assessing the state is too short, we must expect a poor estimate of state, simply because we need to sample long enough to see the fluctuations that define the state. Second, if the true underlying changes in state happen with a typical timescale, say *T*, then we must use an assessment window that is less than *T*. Otherwise, we will be averaging together periods with different states. To understand these constraints quantitatively, we test our approach using surrogate data in which we control exactly when the state changes occur, and we know exactly what the states are (Fig S2a). Our surrogate data switches randomly among three different states, with each state lasting for *T* seconds; we tested a range of *T* between 30 and 300 s. The surrogate data was simulated using three different AR models; one was the best fit model for one of the sleep datasets (red background in Fig S2a), another was the best fit model for wake with low EMG (cyan background in Fig S2a), and the third was the best fit model for wake with high EMG (purple background in Fig S2a). Thus, we know exactly what the true $d_2$ is during each state of the surrogate data and can test how accurately our sliding window measure of $d_2$ recovers the true $d_2$ (Fig S2b). Here accuracy was assessed using two $d_2$ thresholds $\Theta_1$ and $\Theta_2$. If an estimated $d_2 > \Theta_2$, then it was classified as the state with largest ground-truth $d_2$ (sleep). If $\Theta_1 > d_2 > \Theta_2$, it was classified as the state with intermediate ground-truth $d_2$. If an estimated $d_2 < \Theta_1$, then it was classified as the state with low ground-truth $d_2$. All possible thresholds with $\Theta_1 < \Theta_2$ were tried. The accuracy shown in Fig S2b is the fraction of correctly classified $d_2$ estimates for the best choice of $\Theta_1$ and $\Theta_2$. We found good accuracy when we used a 60 s sliding window for assessing $d_2$, for a wide range of state durations. This motivated our decision to use a 60 s sliding window in the main text results. Note that chance level accuracy is 33% for our surrogate data. Finally, we directly tested the robustness of our observed correlations between $d_2$ and EMG (Fig 2f) for a range of moving window durations. For the wake state, we found that the correlation between EMG and $d_2$ was robust for a wide range of moving window durations (Fig S2c), suggesting that EMG-related changes in state are occurring at long time scales in the awake state.

**Determining basins of attraction of each fixed point**

Upon coarse-graining, an AR model will flow to the $\beta$ fixed point if the model lies within the "basin of attraction" of that fixed point. Here we mathematically derive the geometry of these basins of attraction. Let the Taylor expansion of $\tilde{\phi}(\omega)$ around $\omega = 0$ be $\tilde{\phi}(\omega) = \sum_m b_m \omega^m$. Then

$$g(\omega) \propto \left|1 - \tilde{\phi}(\omega)\right|^2 = (1 - b_0)^2 + \mathcal{O}(\omega^2)$$

so the dominant term in $g(\omega)$ is the constant term (and hence the AR model flows into the white noise fixed point) if $b_0 \neq 1$. If instead $b_0 = 1$, then

$$g(\omega) \propto (-b_2 - b_2^* + b_0 b_2^* + b_2 b_0^* + |b_1|^2)\omega^2 + \mathcal{O}(\omega^4) = |b_1|^2 \omega^2 + \mathcal{O}(\omega^4)$$

If $b_1 \neq 0$, then the dominant term in $g(\omega)$ is the quadratic term, so the AR model flows into the $\beta = 2$ fixed point. This is the base case of an induction argument for the following claim: for all $\beta \geq 2$ even, the dominant term in $g(\omega)$ is proportional to $\omega^\beta$ if and only if $b_0 = 1$ and $b_1 = \cdots = b_{\beta/2-1} = 0 \neq b_{\beta/2}$. The base case is $\beta = 2$.

For the induction hypothesis, suppose that the statement holds for $\beta = k$ with $k \geq 2$ even. We want to show that the statement holds for $\beta = k + 2$. If the dominant term in $g(\omega)$ is $\omega^k$, then

$$g(\omega) \propto \left( -b_k - b_k^* + \sum_{j=0}^{k} b_j b_{k-j}^* \right) \omega^k + \mathcal{O}(\omega^{k+2})$$

By the induction hypothesis, $b_0 = 1$ and $b_1 = \cdots = b_{k/2-1} = 0$, so this reduces to

$$g(\omega) \propto \left( -b_k - b_k^* + \left| b_{k/2} \right|^2 + b_0 b_k^* + b_k b_0^* \right) \omega^k + \mathcal{O}(\omega^{k+2}) = \left| b_{k/2} \right|^2 \omega^k + \mathcal{O}(\omega^{k+2})$$

Hence the order $k$ term is zero only when $b_{k/2} = 0$. If the order $k$ term is zero, then the same calculation tells us that the order $k + 2$ term is nonzero only when $b_{(k+2)/2} \neq 0$. The result is that the statement holds for $\beta = k + 2$. We conclude by induction that it holds for all even $\beta \geq 2$. Next observe that

$$\tilde{\phi}(\omega) = \sum_t \phi_t e^{-i\omega t} = \sum_{m=0}^{\infty} \frac{(-i)^m}{m!} \left( \sum_t \phi_t t^m \right) \omega^m$$

so $b_m = \frac{(-i)^m a_m}{m!}$, where $a_m = \sum_t \phi_t t^m$. We conclude that an AR model flows into the $\beta$ fixed point if and only if $a_0 = 1$ and $a_1 = \cdots = a_{\beta/2-1} = 0 \neq a_{\beta/2}$. Let $A_\beta^{(p)}$ be the set of AR(p) history kernels that satisfy these constraints (i.e., the basin of attraction of the $\beta$ fixed point) and let

$$\hat{A}_\beta^{(p)} = \{ \boldsymbol{\phi} \in \mathbb{R}^p : a_0 = 1, a_1 = \cdots = a_{\beta/2-1} = 0 \}$$

Then $\hat{A}_\beta^{(p)}$ is an affine subspace of $\mathbb{R}^p$ with dimension $\max\{0, p - \beta/2\}$ and $A_\beta^{(p)} = \hat{A}_\beta^{(p)} \setminus \hat{A}_{\beta+2}^{(p)}$. The nested structure of the constraints translates to $\hat{A}_{\beta_2}^{(p)} \subseteq \hat{A}_{\beta_1}^{(p)}$ for all $\beta_2 \geq \beta_1$. For example, in the AR(2) case, $\hat{A}_2^{(2)}$ is the line through $(0, 1)$ and $(1, 0)$, $\hat{A}_4^{(2)}$ is the singleton $\{(2, -1)\}$, and $A_2^{(2)}$ (the set of AR(2) models that flow into the $\beta = 2$ fixed point) is the line $\hat{A}_2^{(2)}$ minus the point $\hat{A}_4^{(2)}$.

#### Calculating distances $d_\beta$

Having identified all the basins of attraction, we can now calculate the distances from them to a given AR model (in the main text, this is the AR model obtained from a fit to the experimental data). For a fixed model order $p$ and an AR($p$) model with history kernel $\boldsymbol{\theta} = (\theta_1, \theta_2, \ldots, \theta_p)^T$, define

$$d_\beta = \inf\{ \|\boldsymbol{\theta} - \boldsymbol{\phi}\|_2 : \boldsymbol{\phi} \in A_\beta^{(p)} \}$$

But $\hat{A}_\beta^{(p)}$ is closed, $A_\beta^{(p)} \subseteq \hat{A}_\beta^{(p)}$, and every point in $\hat{A}_\beta^{(p)}$ is realized as the limit of a sequence of points in $A_\beta^{(p)}$, so $\hat{A}_\beta^{(p)} = \mathrm{cl}(A_\beta^{(p)})$ and we can equivalently characterize $d_\beta$ as the distance from $\boldsymbol{\theta}$ to the closest point in $\hat{A}_\beta^{(p)}$.

The first step in calculating $d_\beta$ is showing that, for any $r \geq 1$, the only element $\boldsymbol{\phi}^{(r)}$ of $\hat{A}_{2r}^{(r)}$ is given by

$$\phi_t^{(r)} = \binom{r}{t}(-1)^{t+1}$$

This follows immediately from the binomial theorem, which tells us that for any real numbers $x, y$,

$$(x+y)^r = \sum_{t=0}^{r} \binom{r}{t} x^t y^{r-t} \qquad (S1)$$

Differentiating $m$ times $(1 \leq m \leq r-1)$ with respect to $x$, setting $x = -1, y = 1$, and rearranging terms, we get

$$\sum_{t=1}^{r} t(t-1)\cdots(t-m+1)\phi_t^{(r)} = 0$$

Setting $m = 1$, we find $a_1 = 0$. Plugging this into the equation for $m = 2$ then yields $a_2 = 0$, and so on up to $a_{r-1} = 0$. To show that $a_0 = 1$, we substitute $x = -1, y = 1$ into Eq. (S1):

$$0 = \sum_{t=0}^{r} \binom{r}{t}(-1)^t = 1 - \sum_{t=1}^{r} \phi_t^{(r)} = 1 - a_0 \Rightarrow a_0 = 1$$

Hence $\boldsymbol{\phi}^{(r)} \in \hat{A}_{2r}^{(r)}$. Embedding $\boldsymbol{\phi}^{(r)}$ in $\mathbb{R}^p$ by appending $p - r$ zeros, we have $\boldsymbol{\phi}^{(r)} \in \hat{A}_{2r}^{(p)}$. Now fix an even $\beta \leq 2p$. If $r \geq \beta/2$, by the nested property $\hat{A}_{2r}^{(p)} \subseteq \hat{A}_\beta^{(p)}$, so $\boldsymbol{\phi}^{(r)} \in \hat{A}_\beta^{(p)}$. We conclude that the $p - \beta/2 + 1$ points $\boldsymbol{\phi}^{(\beta/2)}, \boldsymbol{\phi}^{(\beta/2+1)}, \ldots, \boldsymbol{\phi}^{(p)}$ are all in $\hat{A}_\beta^{(p)}$. These are enough to uniquely define $\hat{A}_\beta^{(p)}$ because it has dimension $p - \beta/2$.

Calculating $d_\beta$ now amounts to a well-studied linear algebra problem. Let $X$ be the $p \times (p - \beta/2)$ matrix whose $i$th column is $\boldsymbol{\phi}^{(\beta/2+i)} - \boldsymbol{\phi}^{(\beta/2)}$ for all $i = 1, 2, \ldots, p - \beta/2$. First, we use MATLAB's "qr" function to decompose $X$ into the product of a $p \times p$ orthonormal matrix $Q$ and a $p \times (p - \beta/2)$ upper triangular matrix $R$. We then use MATLAB's "mldivide" function to find the $(p - \beta/2) \times 1$ vector $\boldsymbol{v}$ that minimizes $\left\| R\boldsymbol{v} - Q^T\left(\boldsymbol{\theta} - \boldsymbol{\phi}^{(\beta/2)}\right)\right\|_2$. The closest point in $\hat{A}_\beta^{(p)}$ to the given history kernel $\boldsymbol{\theta}$ is then $\boldsymbol{\phi}^{closest} = X\boldsymbol{v} + \boldsymbol{\phi}^{(\beta/2)}$ and we have $d_\beta = \|\boldsymbol{\theta} - \boldsymbol{\phi}^{closest}\|_2$.
